# Probability-based detection of phosphoproteomic uncertainty reveals rare signaling events driven by oncogenic kinase gene fusion

**DOI:** 10.1101/621961

**Authors:** Xavier Robin, Franziska Voellmy, Jesper Ferkinghoff-Borg, Conor Howard, Tom Altenburg, Mathias Engel, Craig D. Simpson, Gaye Saginc, Simon Koplev, Edda Klipp, James Longden, Rune Linding

**Affiliations:** Biotech Research and Innovation Centre, University of Copenhagen, Copenhagen, DK-2200, Denmark; Institute of Biology, Humboldt Universität zu Berlin, Berlin, 10099, Germany; Mount Sinai Center for Bioinformatics, Icahn School of Medicine at Mount Sinai, New York, 10029-6574, United States of America

## Abstract

We describe a novel Bayesian method for estimating protein concentration and phosphorylation site occupancy ratios from mass spectrometry experiments. Our variance model assigns standard deviations to all quantitative ratios, even when only a single peptide is observed, increasing the number of quantifiable observations in a sample compared to conventional methods. We further demonstrate the application of this method using a dataset investigating the impact of the PRKAR1A-RET gene fusion in immortalized thyroid cells.

## Main text

Typically, the analysis of data generated by mass spectrometry is hampered by few available replicates, therefore harsh filtering applied to ensure statistical significance results in the loss of a substantial amount of data points. This effect is exacerbated when considering analysis of post-translational modifications, which are usually measured as a single peptide, as opposed to proteins that are measured as the aggregate of multiple peptides. Moreover, uncertainties can only be measured reliably when enough measurements are available and are often ignored altogether. In order to provide accurate uncertainties on the measurement of protein and phosphorylation site ratios, we developed a Bayesian noise model of the experimental and instrumental errors, and derived a likelihood expression of protein concentration and phosphorylation site occupancies, sampled using a Bayesian Markov Chain Monte Carlo approach. Through the propagation of effective measurement numbers this model can more accurately derive significance from a low number of observations. In order to model log ratios (sample versus control) of protein concentration (*c*) and occupancy of post-translational modification sites (*o* and *o*′ representing the proportion of proteins where the site is modified in the sample and control, respectively), we established the following formula, which models the average ratio (*μ*) of a peptide *i* (with potentially multiple post-translational modifications) as a function of the parameters *c*, 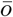 and 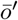:

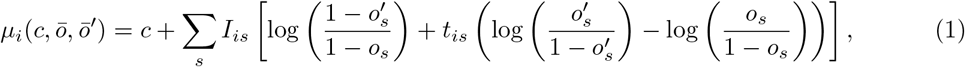

where *I*_*is*_ and *t*_*is*_ are indicator matrices specifying which sites are covered and modified, respectively.

Together with an appropriate noise model, this function could be sampled and compared to experimentally measured peptide ratios with Markov Chain Monte Carlo (Figure 1A-D) resulting in a distribution of likely values for the parameters. To validate the accuracy of our quantitative model we analyzed the UPS1 spike-in standard dataset^1^; a mixture of 48 recombinant proteins spiked at several known concentrations to a fixed background of yeast lysate. After applying normalization, we calculated peptide log ratios to the 12.5 fmol/μg sample, peptide standard deviation and the number of effective observations (Figure 1E). Samples with less than 2.5 fmol/μg were discarded as they contained very few identified UPS1 peptides. We then randomly generated 100 artificial proteins from the remaining peptides with random Gaussian average ratios and sampled these with the Monte Carlo likelihood model, the results of which are shown in Figure 1F. The model produced relatively conservative estimates of the true ratios, and ratios with large uncertainties were drawn towards 0. Overall the correlation of known to predicted concentration ratios was 0.938 validating the accuracy of our method. To further validate our model we reanalyzed previously published phosphoproteomic analyses investigating epidermal growth factor stimulation^2^ and insulin signaling^3^. In all cases the Monte Carlo model was able to quantify, with measures of uncertainty, more proteins and phosphorylation sites than conventional methods, with the most significant increases seen in sparse data with few replicates (Figure S1).

**Figure 1.**
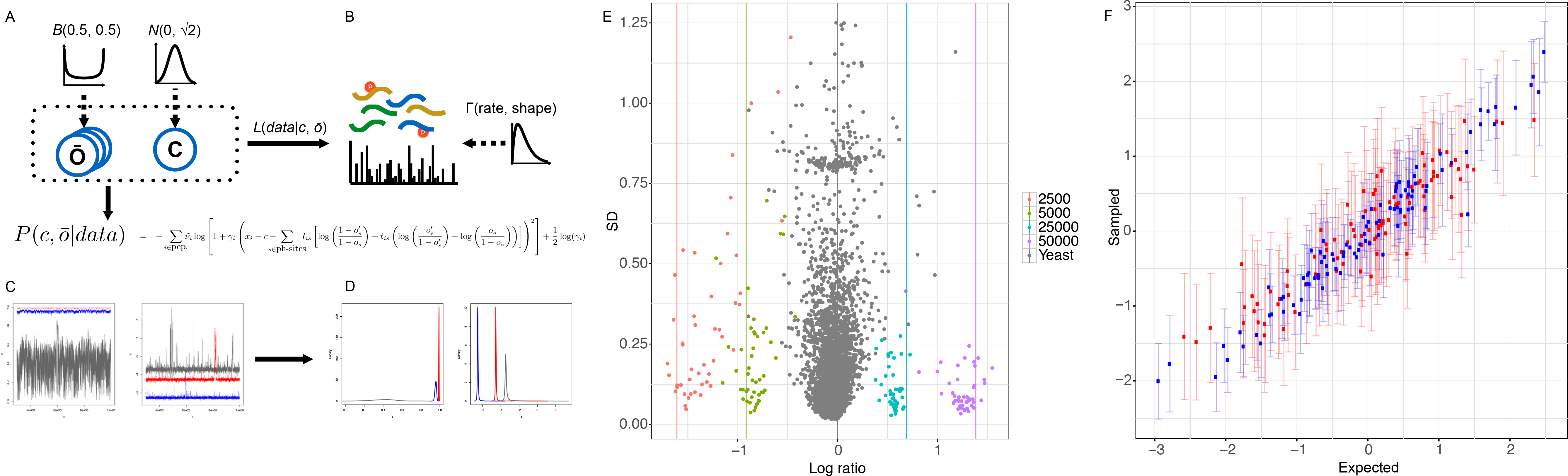
Using the likelihood function and minimum information priors (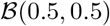 and 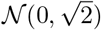), the parameters of interest for phosphosite occupancies 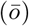 and concentration (*c*) (**A**) could be compared to the data (**B**) and noise model Γ(*rate, shape*). This allowed the expression of the probability of the parameters, given the data 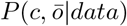, which can then be sampled with Markov-Chain Monte Carlo (**C**), and finally converted to probability distributions (**D**), represented by mean and standard deviations. (**E**) illustrates the observed log concentration ratios for an example dataset of spiked proteins (dots) along with the expected ratio (vertical lines). (**F**) illustrates the correlation observed between expected log ratios of protein concentration and the sampled log ratio following Markov-Chain Monte Carlo modelling for an example dataset of spiked proteins sampled with 1 (red) or 3 (blue) replicates.

In order to demonstrate the applicability of this method for analyzing complex large-scale data, we investigated the effects of the PRKAR1A-RET chromosomal translocation in immortalized thyroid cells using global phospho-proteomics. Fusion kinases are commonly observed in both blood and solid tumors in cancer patients, and many have been shown to be recurrent^4^. The best known of these is the BCR-ABL fusion, a driver for chronic myeloid leukemia, which led to the development of the first targeted therapy for cancer: Imatinib/Gleevac^4^. Given that gene fusions have the potential to be relevant therapeutic targets it is perhaps surprising that the underlying signaling network alterations driving cancer progression are still only described, to some degree, for a handful of gene fusions. It would seem likely that mass spectrometry analysis of fusion-induced signaling network rearrangements could go some way towards filling this void. However, a limitation to such studies is that often peptides ‘fall out’ of the analysis due to limitations of instrumentation and sample preparation. Thus, data analytical methods that can help capture low-stoichiometry events are critically needed.

The PRKAR1A-RET fusion kinase has been observed in multiple thyroid cancer patients retaining the regulatory and cyclic NMP binding domains of the PRKAR1A protein on the N-terminus and the full tyrosine kinase domain from the RET kinase on the C-terminus^5^. In order to create the most physiologically relevant model we recreated this fusion using CRISPR-Cas9 in thyroid cells. As primary human epithelial cells senesce after only a few divisions in culture we used SV-40 immortalized thyroid cells (Nthy-ori 3-1^6^) that are suitable for large-scale mass spectrometry experiments, but have low colony formation ability and are non-tumorigenic in mice. Following expression of the PRKAR1A-RET fusion, cells were tested in an anchorage-independent growth assay. Strikingly, we observed a significant increase in colony formation in the cells expressing the PRKAR1A-RET rearrangement compared to control cells (Figure S1). This would suggest that the engineered fusion event, either directly or indirectly, induced malignant transformation in our cell model. Mass spectrometry analysis of the cells and colonies expressing the PRKAR1A-RET gene fusion identified 1985159 peptides, 248536 of which were phosphorylated, over the entire presented dataset. This corresponded to 5759 unique proteins and 11435 unique, high-confidence, phosphorylation sites; 5672 of the unique proteins were observed with two or more peptides. From these data our Monte Carlo model was able to quantify ratios, calculated from sample thyroid cells expressing the PRKAR1A-RET fusion to control thyroid cells expressing Cas9 protein alone, for 94% of the observed proteins (5411) and 70% of the observed phosphorylation sites (8032), significantly more than was possible using conventional analysis methods (MaxQuant).

We noted that a remarkably similar phosphorylation site occupancy signal was observed in each fusion-expressing cell line (Figure 2), suggesting that the changes to signaling networks driven by this chromosomal rearrangement are highly conserved. Given that the cells grown in colony formation assays were also distinct from fusion-kinase expressing cells at the level of both the proteome and phospho-proteome this would suggest that additional signaling network rearrangements occurred driving tumorigenesis in cells that displayed anchorage-independent growth. In particular we noted that the concentration of PRKAR1A was found to be downregulated in the transformed cells relative to colonies formed without the fusion construct. In order to determine which cellular processes were regulated following PRKAR1A-RET fusion functional annotations were assigned using the hallmark gene set collection of the Molecular Signature Database^7^, based on coherently expressed genes representative of biological processes^8^. Processes that were significantly enriched in cells expressing the PRKAR1A-RET fusion are shown in Figure S2 and Table S1. In particular, we noted that in cells expressing the PRKAR1A-RET fusion there was enrichment for signaling via PI3K-AKT-mTOR, which has previously been shown to affect cell proliferation, viability and iodide uptake in thyroid cells^9^, and Myc, which is known to be involved in the majority of human cancers and leads to the acquisition of hallmark cancer characteristics such as uncontrolled proliferation^10^. Interestingly we observed that additional signaling pathways were modulated in the colonies expressing PRKAR1A-RET, including P53, alterations in which are known to contribute to carcinogenesis in thyroid tissue^11^. Given the significant number of additional phosphorylation sites quantified by the Monte Carlo model (most likely due to the sparcity of the data, which has fewer biological replicates than the other analyzed datasets) we used DAVID^12^ to determine whether there was enrichment of specific signaling pathways which were only quantified by Monte Carlo analysis. We observed significant enrichment of proteins associated with spliceosome (P=0.0026) and glycolysis (P=0.018) pathways, both of which are considered relevant targets for novel cancer therapeutics^13,14^. Given this, and the known importance of kinases in cancer^15^ in general, it would seem highly likely that the additional information provided by the Monte Carlo model would aid biological analysis of the role of the PRKAR1A-RET fusion kinase.

**Figure 2.**
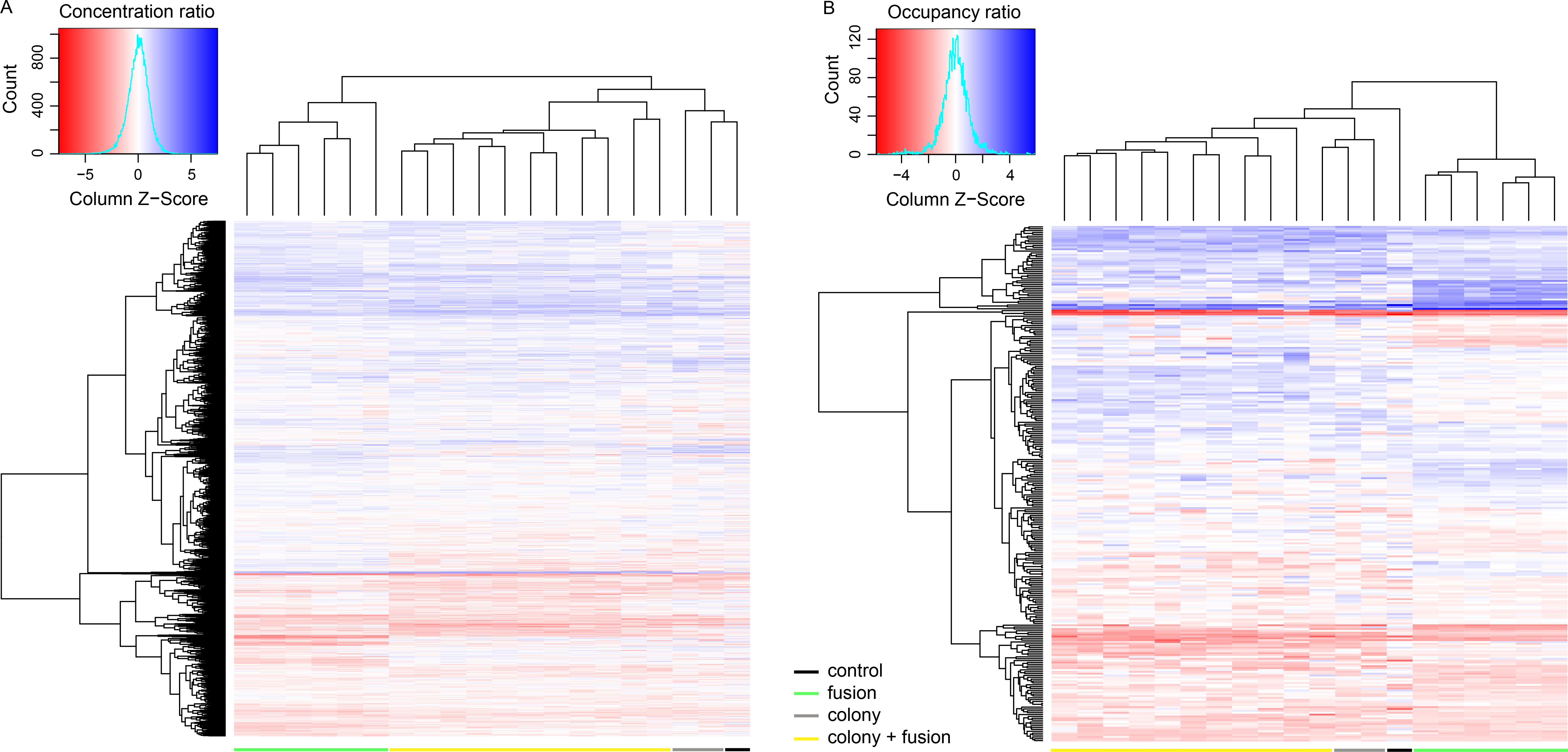
Hierarchical clustering of Markov-Chain Monte Carlo modelled data using a Euclidian distance metric distinctly clustered the fusion-kinase expressing cells from control cells, and fusion-kinase expressing cells grown in standard cell culture from those grown in colony formation assays. The concentration of quantified proteins (**A**), and the occupancy of quantified phosphorylation sites (**B**) were expressed as log ratios to thyroid cells expressing Cas9.

These data demonstrate the ability of our model to estimate protein concentration and phosphosite occupancies and to provide an accurate estimate of the uncertainty of those parameters. We believe that a more rigorous modeling of uncertainties is a critical step towards more precise models, which will allow more informed interpretation of quantitative mass spectrometry data. While the model presented here is based on peptide ratios, this method could also be applied to model peptide intensities. This would enable the modeling of additional missing data points due to intensities that are below the detection threshold of the mass spectrometer. Similar models could be used for datasets generated using other quantification methods such as TMT or SILAC. Moving forward we expect that likelihood-based algorithms taking full advantage of measurement uncertainties could aid in the design of new mass-spectrometers by searching for bottlenecks and limitations in the ion path of the instruments. In addition, we have demonstrated the application of this model to a study sampling the proteomic and phospho-proteomic network changes upon endogenous expression of a chimeric gene found in solid tumors. This analysis confirms the regulation of proteins known to be involved in the progression of thyroid cancers. Furthermore, our analysis shows that cells harboring the induced fusion are enriched for processes essential for tumorigenesis.

## Methods

### Cell lines and vectors

SV40 immortalized primary cells (Nthy-ori 3-1) from thyroid follicular epithelial tissue were purchased from Sigma and cultured in RPMI 1640 medium supplemented with 2 mM l-glutamine, 10% fetal bovine serum and 1% penicillin/streptomycin solution (all ThermoFisher Scientific) in a humidified environment at 37°C with 5% CO_2_. HEK-293T cells were cultured in Dulbecco’s Modified Eagle Medium, supplemented with 10% fetal bovine serum and 1% penicillin/streptomycin. pCMV-VSV-G, psPAX2, pCW-Cas9 Dox-inducible lentiviral vector and pLX-sgRNA lentiviral vector containing AAVS1-targeting sgRNA were all obtained from Addgene.

### Determination of targeted breakpoint location

The empirically observed breakpoint of the translocation event was verified using COSMIC^16^ and ChiTars^17^ and the introns selected by following the principle of last observed- and first observed exons given several documented cases of the same translocation, with varying numbers of exons detected in the partner genes. Only one combination of introns targeted for CRISPR-Cas9-mediated cleavage was retained, using the intron following the last seen exon of the head partner and the intron preceding the first observed exon of the tail moiety.

### CRISPR sgRNA target selection and cloning

All SpCas9 Protospacer Adjacent Motif (PAM) sites within selected introns were scored using the method described by Doench et al.^18^. The two top-scoring guide sequences were retained for each translocation partner. A ‘G’ was prepended to the guide sequence (GX20 sgRNA), with the benefit of diminishing off-targets at the cost of cutting efficiency^18^. Top-scoring guide sequences were verified by submission to the CRISPR Design webtool (http://crispr.mit.edu/)^19^. Selected sgRNAs were verified with the updated off-target prediction algorithm GUIDE-seq^20^. The pLX-sgRNA vector was digested with Xba1, Nde1 for 2 hrs at 37°C, ran on a 0.5% agarose gel followed by gel extraction and purification. The DNA digest was further used to clone a new vector allowing for facile insertion of sgRNA oligonucleotides via Gibson reactions. In order to select for successful transduction of the second sgRNA, the vector was further modified to replace the blasticidin resistance with hygromycin.

### HEK-293 infection

2×10^6^ HEK-293T packaging cells were seeded in a 75cm^2^ cell culture flask in growth medium without antibiotics. The following day, cells were transfected with pCMV-VSV-G, psPAX2, and pCW-Cas9 vectors using MirusBio TransIT-LT1 transfection reagent. After 24 hours medium was removed and fresh medium containing 10mg/ml bovine serum albumin was added. 24 hours later medium was harvested and residual cells were removed by filtration.

### Viral pCW-Cas9 transduction

Nthy-ori 3-1 cells were seeded at 300 000 cells/well in a 6-well plate. The following day, protamine sulfate was added to flasks to a final concentration of 8 *μ*g/ml and cells were transduced with the Cas9 construct with increasing volumes of virus-containing harvested medium: 0, 2, 5, 10, 20, 50, 75, 100, 150, 200, 250 *μ*L. After overnight incubation, growth medium was refreshed. Cells with increasing viral titers were seeded in 96-well tissue culture plates at 5000 and 10000 cells/well in 200 *μ*L of media. Growth media was replaced with medium supplemented with 0.8 *μ*g/mL puromycin the following morning. Alamar Blue was added to the wells and the percent viable cells were measured at 570 nm after 5 hours and 24 hours. The lowest viral titer at which cells were viable post selection was chosen, and cells were further maintained in puromycin-containing media.

### Viral modified pLX-sgRNA transduction

Medium containing viral sgRNA vectors was harvested as previously described for each individual sgRNA. 1×10^6^ Thy-Cas9 cells were seeded per 25 cm^2^ cell culture plate. The following morning harvested medium was added to growth medium in increasing volumes, at equal volumes between the two sgRNAs of a given combination. Cells were split twice and selected for 14 days in medium supplemented with 0.8 *μ*g/mL puromycin, 10 *μ*g/mL blasticidin or 200 *μ*g/mL hygromycin. The condition with lowest viral volume mediating resistance was retained for further experiments.

### Cas9 expression induction

Doxycycline at a final concentration of 1 *μ*g/mL was administered for 1 week to 4×10^5^ cells seeded in a 100mm tissue culture dish. Doxycycline-containing medium was refreshed after 4 days. Immediately thereafter individual clones were generated via single cell cloning and expanded.

### Translocation detection PCR

RNA was extracted from cells at various passages following single cell cloning using a Qiagen RNA extraction kit according to the manufacturers instructions. Clones were screened for translocations and the cDNA breakpoint determined by RT-PCR in a 96-well AB Veriti Fast Thermal Cycler using the Quantitect SYBR green RT-PCR kit (Qiagen) according to the manufacturers instructions with an annealing temperature of 56°C for 30 seconds, an extension at 72°C for 60 seconds, and a final extension for 5 minutes at 72°C. The forward primer was AGACAATGGCCGCTTTAGCCA and the reverse primer was CAGGGAGCCGTATTTGGCGT. Sanger sequencing was used to confirm the sequence of bands at the expected weight.

### Anchorage-independent growth assay

Transformation was verified using a soft agar colony formation assay^21^. 6-well tissue culture plates were coated with growth medium containing 1% agarose, and triplicate dishes were inoculated with 10 000 or 2000 cells suspended in RPMI-1640 medium containing 0.8% agarose and 15% fetal bovine serum, for counting and colony picking, respectively. Colony formation was assessed after 2 weeks. Image analysis was performed using ImageJ, with plugins ColonyArea^22^ and ColonyCounter (https://imagej.nih.gov/ij/plugins/colony-counter.html).

### Mass spectrometry lysis and sample preparation

Biological replicates were generated by selecting several fusion-containing cell lines as well as several cell lines following colony formation. Protein was harvested by lysing a 150 mm dish at 80-90% confluency in 1 mL of modified RIPA buffer (50 mM Tris pH 7.5, 150 mM NaCl, 1% NP-40, 0.5% Na-deoxycholate, 1 mM EDTA) supplemented with phosphatase inhibitors (5 mM b-glycerophosphate, 5 mM NaF, 1 mM Na3VO4 and protease inhibitors (Roche cOmplete ULTRA Tablets, EDTA-free, #05892791001), sonicated, and centrifuged for 20 minutes at 4300G. Ice-cold acetone was added to the supernatant to achieve a final concentration of 80% acetone, and protein was left to precipitate overnight at −20°C. Precipitated protein was pelleted by centrifugation at 1800G for 5 minutes and solubilized in 6 M urea, 2 M thiourea, 10 mM HEPES pH 8.0. Protein was quantified using the Bradford assay and 650 *μ*g of each sample were reduced with 1 mM DTT, alkylated with 5 mM ClAA, and digested with the endopeptidase Lys-C (1:200 v/v, Wako #129-02541) for 3 hours. Samples were diluted to a 1 mg/mL protein concentration using 50 mM ammonium bicarbonate and incubated overnight with trypsin (1:200 v/v, Sigma #T6567). Digested samples were acidified and urea removed using a Sep-Pak C18 96-well plate (Waters #186002320). An aliquot of 50 *μ*g of eluted peptides was set aside for proteome analysis as well as peptide quantitation (using a Pierce quantitative colorimetric peptide assay, Thermo #23275) in order to equalize injection load for label-free LC-MS/MS analysis. The remainder was lyophilized, aliquots of 200 *μ*g made up in 1 M glycolic acid, 80% ACN, 5% TFA, and enriched for phosphopeptides using MagReSyn Ti-IMAC beads (#MR-TIM010 ReSyn Biosciences) according to the manufacturers instructions. Enriched peptides were subjected to a final C18 clean-up prior to data acquisition. All samples were processed as technical duplicates.

### MS data acquisition

All spectra were acquired on an Orbitrap Fusion Tribrid mass spectrometer (Thermo Scientific) operated in data-dependent mode coupled to an EASY-nLC 1200 (Thermo Fisher Scientific) liquid chromatography pump and separated on a 50cm reversed phase column (Thermo, PepMap RSLC C18, 2uM, 100 AA, 75um×50cm). Proteome samples (non-enriched) were eluted over a linear gradient ranging from 0-11% ACN over 70 min, 11-20% ACN for 80 min, 21-30% ACN for 50 min, 31-48% ACN for 30 min, followed by 76% ACN for the final 10 min with a flow rate of 250 nL/min. Phosphopeptide-enriched samples were eluted over a linear gradient ranging from 0-18% ACN over 195 min, 18-26% ACN for 30min, 26-76% ACN for 10min, followed by 76% ACN for the final 5 min with a flow rate of 250 nL/min. Survey full scan MS spectra were acquired in the Orbitrap at a resolution of 120 000 from m/z 350-2000, AGC target of 4×10^5^ ions, and maximum injection time of 20 ms. Precursors were filtered based on a charge state of 2 and monoisotopic peak assignment, and dynamic exclusion was applied for 45 seconds. A decision tree method allowed fragmentation for ITMS2 via ETD or HCD, depending on charge state and m/z. Precursor ions were isolated with the quadrupole set to an isolation width of 1.6 m/z. MS2 spectra fragmented by ETD and HCD (35% collision energy) were acquired in the ion trap with an AGC target of 1×10^4^. Maximum injection time for HCD and ETD was 200 ms for phosphopeptide-enriched samples, and 80ms for proteome samples.

### Fasta file generation

DNA was extracted from the parental cell line using the Qiagen QIAamp DNA Mini kit, according to the manufacturers instructions. Whole exome sequencing was performed by GENEWIZ with a target average exome coverage of 89x. Reads were filtered using Trimmomatic^23^ with the following parameters: headcrop = 3, minlen = 30, trailing = 3. Trimmed reads were aligned to the hg19 reference genome using Burrows-Wheeler transform^24^. The Genome Analysis Toolkit base quality score recalibration was applied^25^ along with indel realignment, duplicate removal, and SNP and INDEL discovery and genotyping across all samples simultaneously using standard hard filtering parameters according to Genome Analysis Toolkit best practice recommendations^26^. The Ensembl Variant Effect Predictor^27^ with Ensembl v. 88 was used to predict the effect of the mutations on the protein sequence. In order to find non-reference, mutated peptides in the mass spectrometry data, we increased the search Fasta file with mutations affecting the protein sequence that passed a high sensitivity filter: QD ≤1.5, FS ≥ 60, MQ ≥ 40, MQRankSum ≤ −12.5, ReadPosRankSum ≤ −8.0, and DP at least 5 per sample on average.

### Spectral search and quantification

All label-free raw data were searched using MaxQuant 1.5.1.2^28^ against the human (or Mouse in the case of Humphrey et al.) Ensembl database (v88), for the PRKAR1A-RET fusion kinase dataset the mutant proteins identified by whole exome sequencing as well as the sequence of the fusion product were also included. Default settings were used with the exception of a minimum peptide length of 6 amino acids. STY phosphorylation was added as a variable modification. An FDR of 0.01 was applied at the level of proteins, peptides and modifications. Phosphosites were filtered for a localization score of at least 0.9.

### UPS1 benchmark

Data was downloaded from ProteomeXchange (PXD001819) and searched with MaxQuant 1.5.1.2^28^ against the yeast protein database from Ensembl 89 and the UPS spike-in Fasta file obtained from Sigma-Aldrich.

### Post-spectral search data processing

Mass spectrometry spectra were identified by MaxQuant^28^ and assigned a sequence and modification status. This was then used to identify the quantitative information obtained from the sequence of MS^1^ spectra over chromatography time, in the form of an Extracted Ion Chromatogram (XIC) measurement of intensity. We then investigated the XIC intensity of a modified peptide with a known sequence, in the corresponding sample and mass spectrometry run (and raw file). We extracted quantitative information regarding ion intensities from the evidence.txt file. We filtered out peptides that were associated with multiple identifications in the msms.txt file, had a score < 40, were identified in the reverse database or came from known contaminants. We also filtered all modified peptides with a localization probability < 0.9.

### Calculating average intensity

A peptide *P* could be observed in multiple XIC in a single run. We defined the aggregated intensity of peptide P_p,s,r_, where *p* was the sequence of the peptide, *s* was the sample and *r* was the run, as:

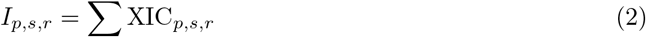

When we observed the peptide in multiple replicates we further summarized the mean intensity I:

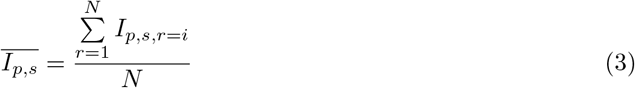

For normalization purposes we also calculated an average intensity over a mass spectrometry run. However because intensities are not normally distributed but approximately follow a log-normal distribution, the log intensities were summarized instead thusly:

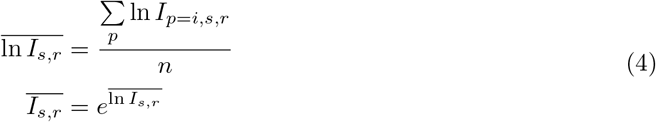

### Ratios calculation

We defined the log-ratio of a peptide concentration from a sample *s* to the control *s* = *u* as:

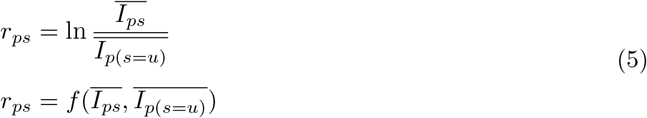

where *I*_*p*(*s*=*u*)_ is the intensity of a peptide *p* in the control sample *u*.

### Error propagation

In order to propagate the standard deviation *σ*_*x*_, when calculating a log-ratio by a pair of intensities we applied the general Gaussian error propagation, which reads for a function f(x,y) depending on the two variables x and y:

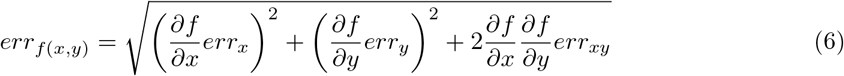

and subsequently in our case for the log-ratio (where the minus is introduced by the partial derivative of the log-ratio to the denominator):

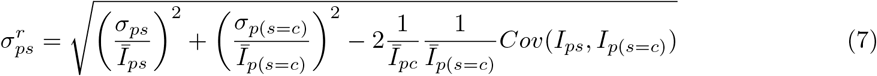

Because we considered the replicates to be independent from each other, we could further simplify the covariance term:

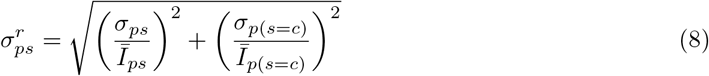

### Normalization

As different amounts of sample could be injected into the mass spectrometer, and because the chromatography is not expected to be identical in each experiment, different experiments were scaled before they were compared using a correction factor based on the average intensity (geometric mean) in order to account for the potential impact of very high intensity peptides within a sample:

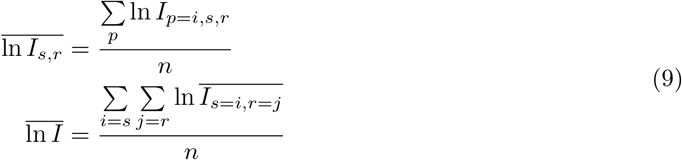

And the correction factor *S* could be expressed as:

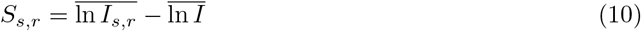

Peptide intensities could thus be corrected, and a normalized intensity *N* calculated, as follows:

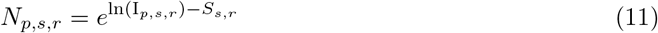

Summarization worked as described above by replacing I_*p,s,r*_ with *N*_*p,s,r*_.

### Variance model

In order to get a baseline estimate of the peptide variance, we established a Bayesian variance model. We modeled the behaviour of the standard deviation of 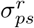 as a function of *abs*(*r*_*ps*_). Data was split into bins containing at least 600 data points each along *abs*(*r*_*p*_*s*), and for each bin we fit 1*/σ*^2^ to a Γ distribution with shape parameter *a* and rate *b*. We observed that *a* ≈ 1, and 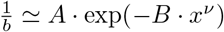, *x* = |log(r)|

### Likelihood function

The likelihood function was calculated from a given phosphorylation site *s*, the set of phosphorylation sites 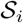 for sequence *i*, the total number of modifiable sites *S* and the occupancy ratio *o*_*s*_ = *p*_*s*_/*c* on site *s*, which is the fraction of peptide that is modified. Assuming *o*_*s*_ values would be mutually independent, the log concentration ratio, *μ*_*i*_, was given by:

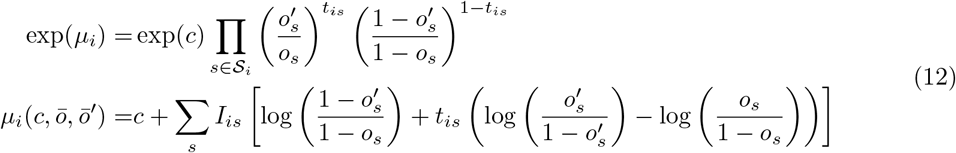

where *c* is the protein log concentration ratio between the sample and the reference, 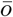 and 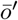 are the collection of occupancy ratios for the sample and the reference respectively, and *t*_*is*_ are indicator variables signifying whether site *s* is modified (1) or not (0). Similarly, *I*_*is*_ = 1 if phospho site *s* was covered by *i* and zero otherwise. 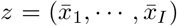 represented the measured log concentration ratios for the *I* peptides belonging to the given protein, where each 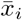 represented the mean value taken over *n*_*i*_ repeats and with effective posterior Γ parameters of *a*_*i*_ and *b*_*i*_ and 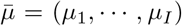 represented the collection of expected values. Then 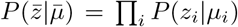 where each factor is a non-standardized *t* distribution. Defined for convenience as:

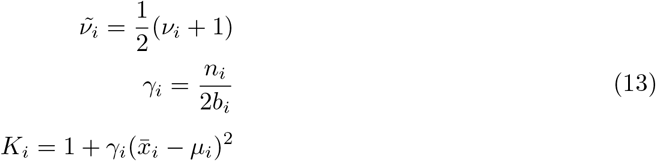

Consequently the likelihood function was given as:

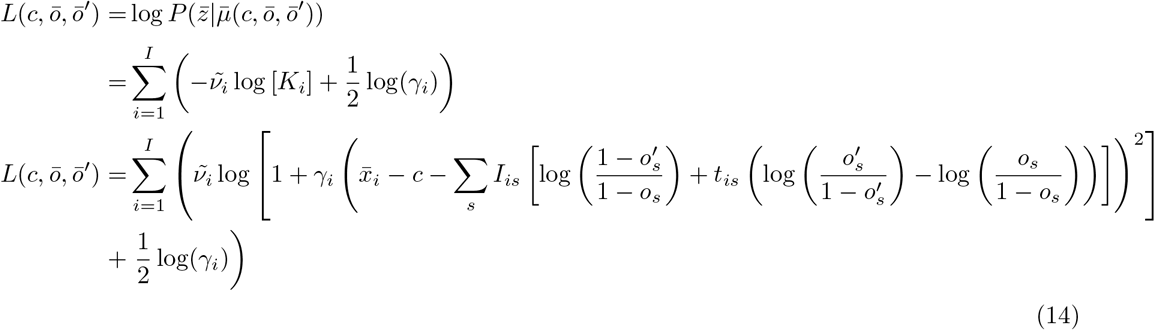

### Markov Chain Monte Carlo sampling

We optimized the function for each protein separately with Bayesian Markov Chain Monte Carlo simulations. We used a Jeffrey prior on *o*_*s*_ with 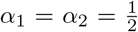 and an exponential prior with *λ* = 2 on *c*.

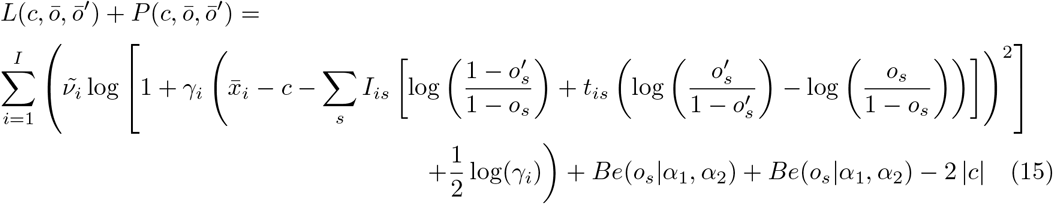

2% of the proposed moves were drawn from the prior distribution of the parameters as described above. Standard moves were as 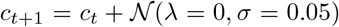 for the concentration and 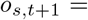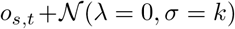 for occupancies, with a standard deviation corrected to propose smaller moves when *os* approaches 0 or 1.

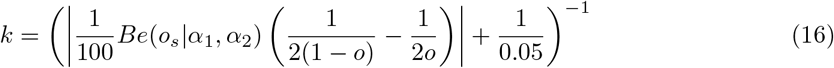

In addition, all moves that would result in *o*_*s*_ < 10^−5^ or *o*_*s*_ > 1 − 10^−5^ were automatically rejected. In order to ensure a good sampling of the likelihood, we used the *effectiveSize* Coda R package^29^ to calculate the effective sample size of the Markov chains. We expressed the number of required iterations as a function of the number of chains *n*_*p*_ as:

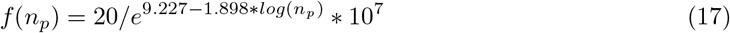

We rounded *f*(*n*_*p*_) up to the next power of 10 to get the final number of iterations *n*_*c*_. Chains with *n*_*c*_ > 10^9^ were re-initialized after 10^9^ iterations. We used a burn-in time of 30% and saved the state of the chains 7 ∗ 10^3^ times as output. Proteins with at least one chain with *effectiveSize* smaller or equal to 100 were repeated with 10 times more iterations.

### Data and software availablility

An R package for performing Markov-Chain Monte Carlo sampling of uncertainties for protein concentration and phosphorylation site occupancy ratios is available at the following URL, under a GNU General Public License Version 3: https://github.com/xrobin/MCMS

## Acknowledgements

This work was funded by the Innovation Fund Denmark Grand Solutions project MorphoMap (1311-00010B) and a Lundbeck Foundation Fellowship both awarded to Rune Linding and the German Research Council (DFG) through the Computational Systems Biology research training group (RTG 1772) awarded to Edda Klipp.

## Author contributions

R.L. and J.F-B. conceived the project. X.R., J.F-B., T.A., M.E. and S.K. created the model. F.V., C.H., C.D.S. and G.S. performed the experiments. X.R. and F.V. analyzed the data. E.K., J.L. and R.L. supervised the research. X.R., F.V., J.L. and R.L. wrote the manuscript.

## Competing interests

The authors report no competing interests.

## Figure legends

Figure S1. Comparison of the number of proteins and phosphorylation sites quantifiable, with uncertainties, using the Monte Carlo model described here and conventional analyses for increasing numbers of replicates.

Figure S2. Soft agar colony formation assays were performed to assess the capacity of PRKAR1ARET kinase fusion expressing cells for anchorage-independent growth. Changes in colony area (**A**) and intensity (**B**) are shown for triplicate repeats. Control cells did not express the PRKAR1A-RET kinase fusion.

Figure S3. Cancer hallmark pathways found to be enriched in PRKAR1A-RET kinase fusion expressing cells following 2D growth (**A**) and anchorage-independent i.e. 3D growth (**B**). Nodes were found to have significantly modulated concentration or phosphorylation site occupancy following Markov-Chain Monte Carlo modelling. Edges were obtained form the STRING database of protein-protein interaction networks^30^.

Table S1. Further processes enriched in PRKAR1A-RET kinase fusion expressing cells and colonies. Only processes with a q-value lower than control cells were included.

