## Supplementary figures and images for "Probability-based detection of phosphoproteomic uncertainty reveals rare signaling events driven by oncogenic kinase gene fusion"

### Supplemental Figure 1

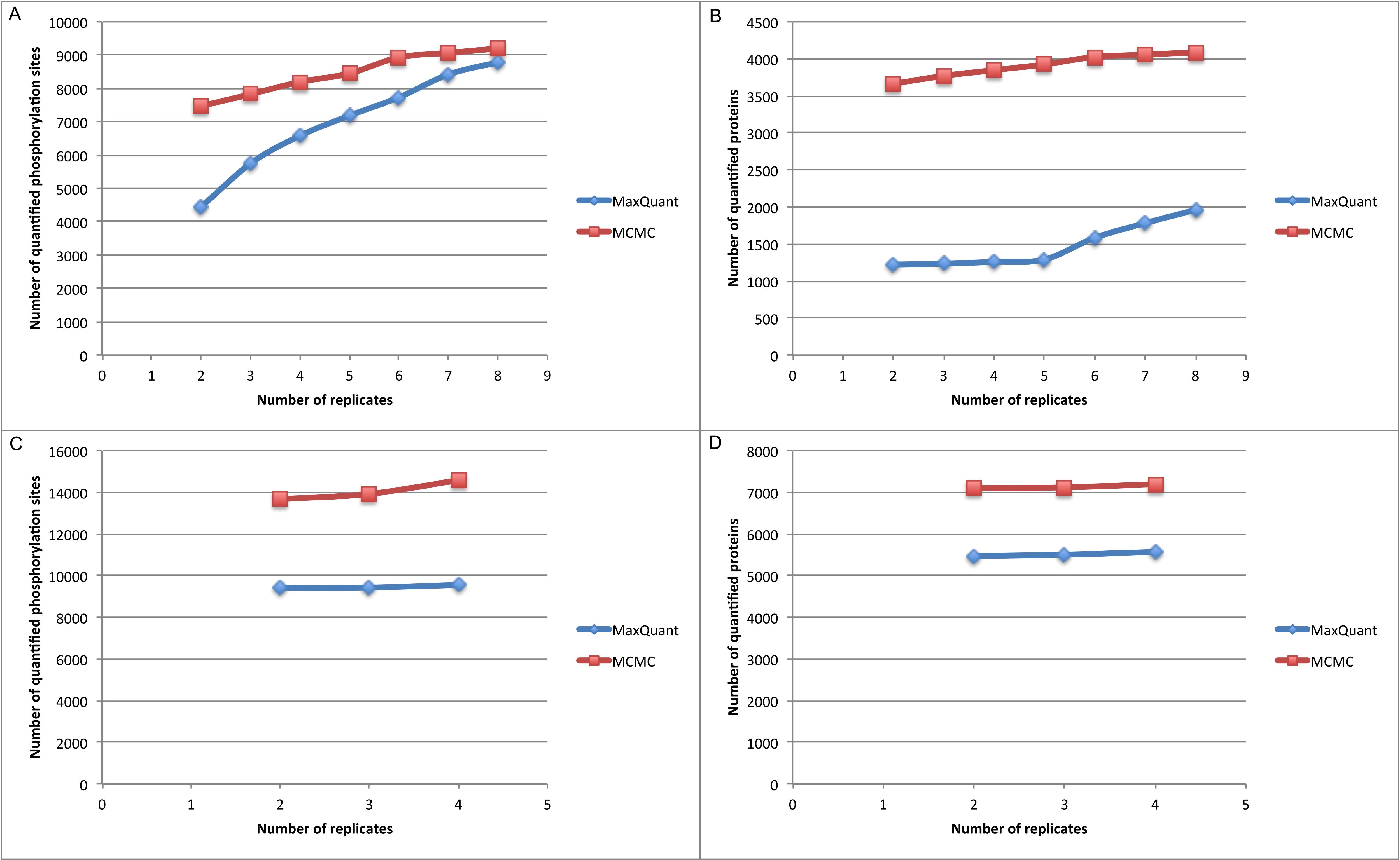

### Supplemental Figure 2

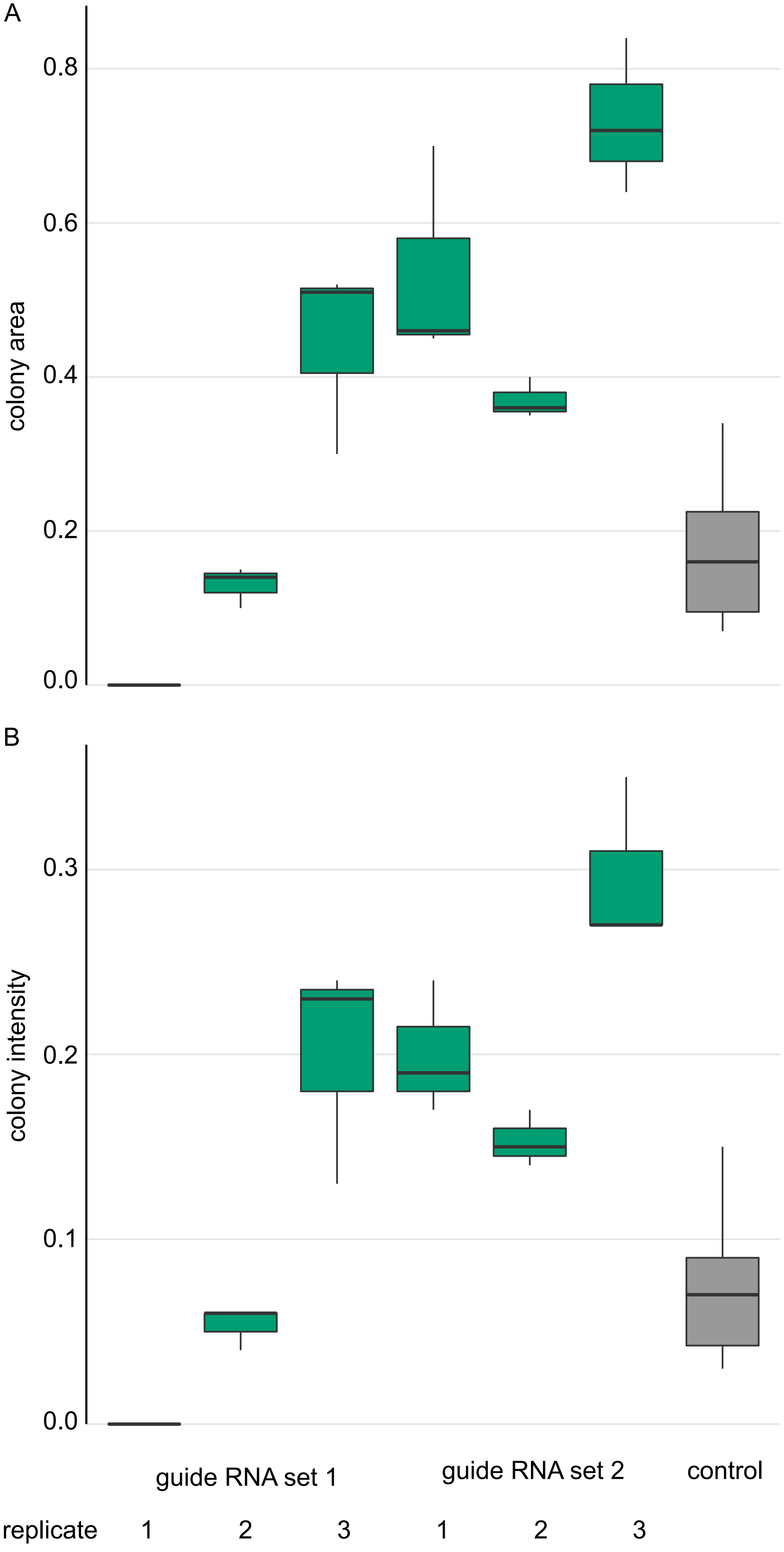

### Supplemental Figure 3

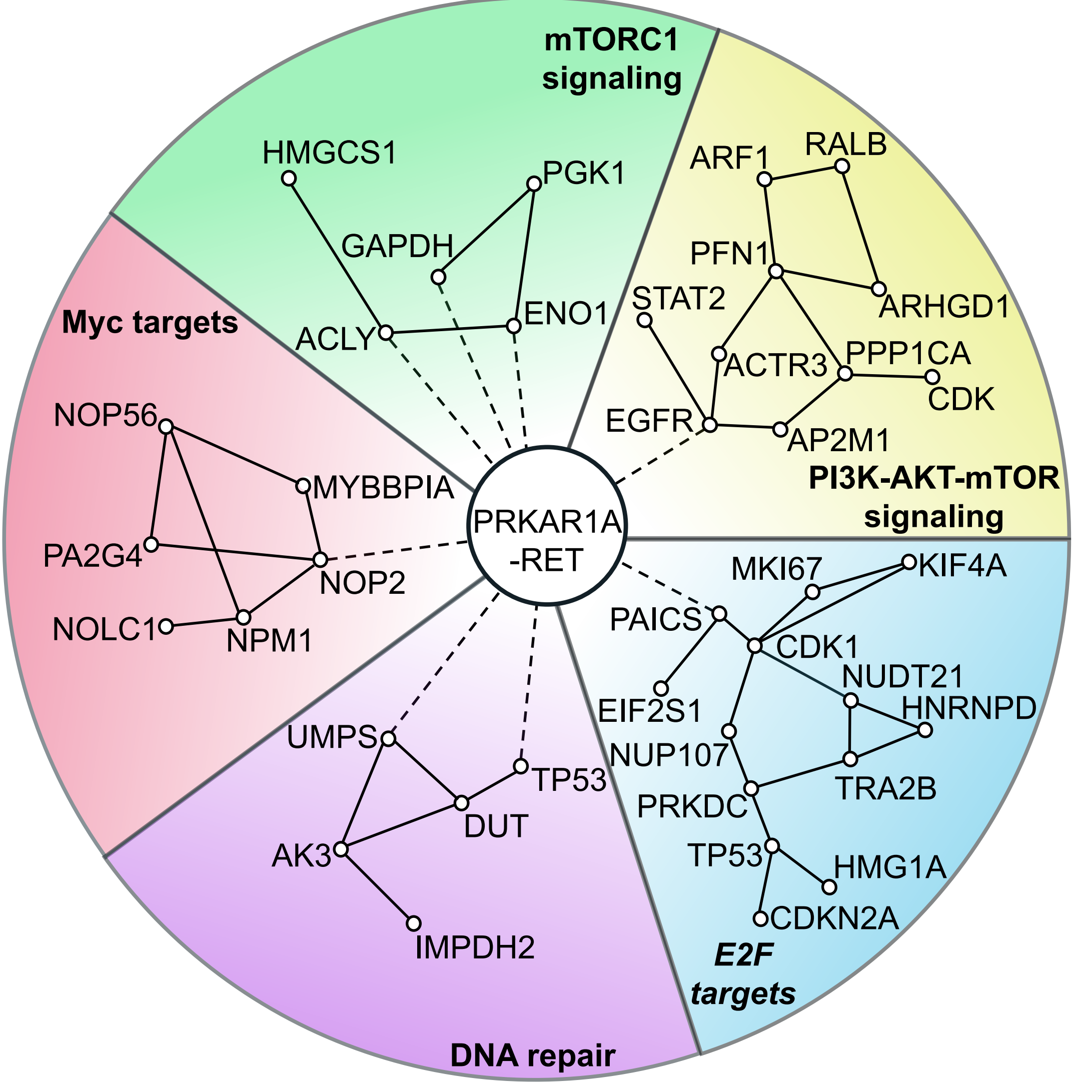

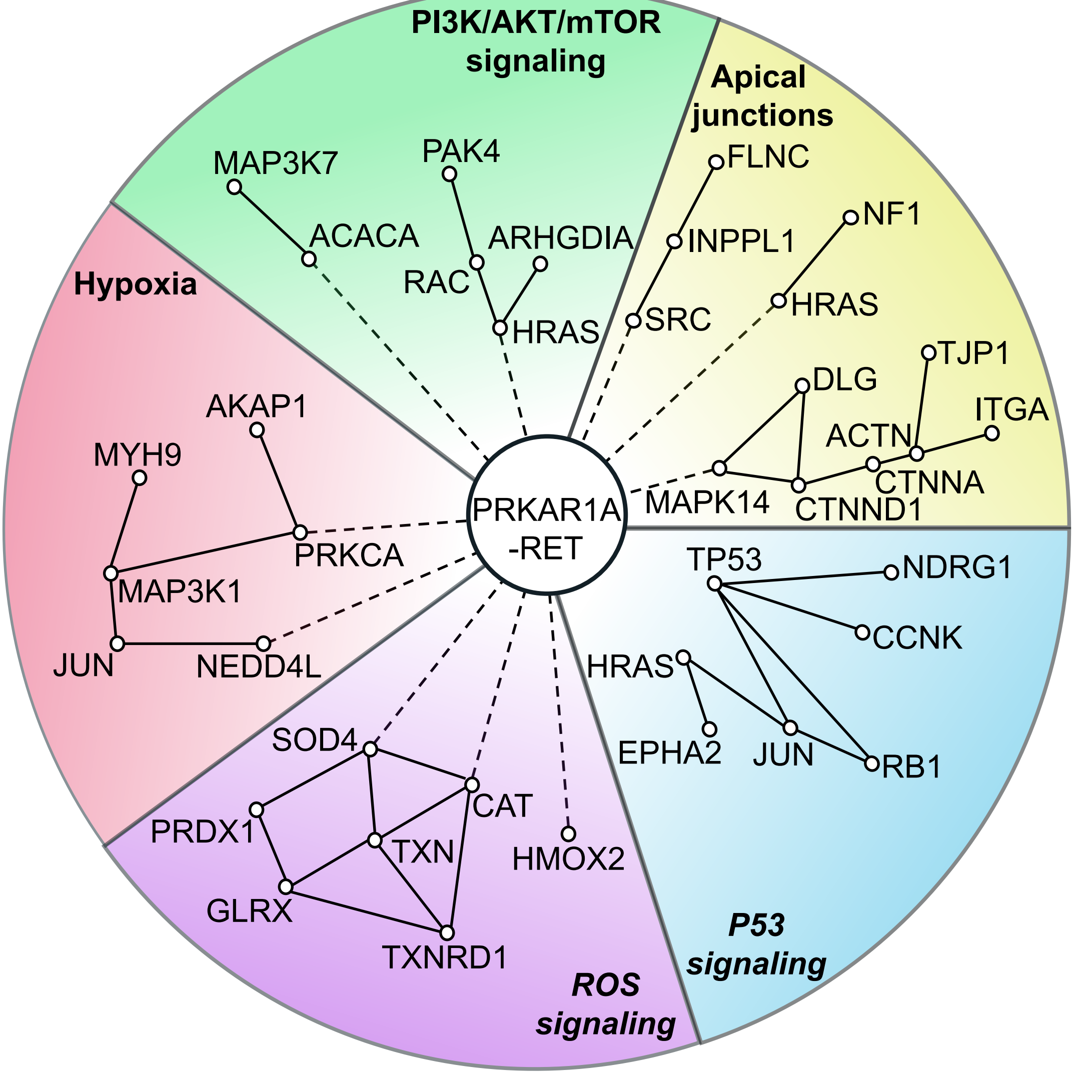
